# Neurophysiological Evidence for Reduced Use of Prior Sound Patterns to Shape Speech Processing in Autism

**DOI:** 10.64898/2026.07.09.737536

**Authors:** Joseph C.Y. Lau, Jacie R. McHaney, Lindsay Goldman, Kylie Robinshaw, Felicia Mou, Kailyn McFarlane, Bharath Chandrasekaran, Molly Losh

## Abstract

Reported perceptual differences in autism may arise from reduced use of prior context to shape incoming sensory input. Speech perception provides a critical test of this account because stable perception requires listeners to integrate variable acoustic signals with contextual expectations. This study examined context-dependent modulation of speech encoding in autistic and non-autistic adults using the frequency-following response (FFR), a neurophysiological measure of phase-locked auditory encoding. Participants heard English intonational pitch contours presented in repetitive and variable contexts while EEG was recorded. Principal component analysis of FFR metrics yielded components indexing neural encoding fidelity and timing. Non-autistic participants showed enhanced encoding fidelity in more predictable contexts, whereas autistic participants showed reduced context-dependent modulation. Neural encoding timing also showed divergent context effects across groups, suggesting altered balance between feedback-based predictive mechanisms and locally driven adaptation processes. Within the autistic group, greater context-related modulation of encoding fidelity was associated with lower ADOS-2 Social Affect severity but poorer speech-in-noise perception, suggesting that the functional impact of contextual modulation depends on input reliability and task demands. These findings indicate that context-dependent modulation of speech encoding is altered in autism and may contribute to individual differences in auditory and social-communicative function.

**Lay Summary:** When we listen to speech, the brain often uses sounds it has recently heard to help make sense of new ones, which is especially useful in noisy, everyday settings. In this study, autistic adults’ brains tended to process speech differently, drawing less on recent sound patterns than non-autistic adults, and this difference was related to their autism domain variability and to understanding speech in background noise. These effects were not simply better or worse but depended on the listening conditions, suggesting that natural differences in how the brain uses recent context may shape some of the ways autistic people listen to speech.

## Introduction

Contemporary theoretical accounts suggest that perceptual differences in autism arise from how prior sensory experience is represented and used to generate expectations about incoming input (Hadad and Yashar, 2022; Lawson et al., 2017, 2014; Palmer et al., 2017; Pellicano and Burr, 2012; Sinha et al., 2014). Within Bayesian (Palmer et al., 2017; Pellicano and Burr, 2012) and predictive processing frameworks (Sinha et al., 2014), perception is an inferential process in which bottom-up sensory signals are integrated with top-down predictions derived from prior context and continuously updated to enable efficient interpretation of dynamic sensory environments. Autistic perception may be characterized by less precise or less strongly weighted priors, resulting in reduced contextual modulation and greater reliance on bottom-up sensory input.

Speech perception is a core function that critically depends on integrating sensory input with multiple sources of contextual information. Speech signals are highly variable and lack a one-to-one mapping onto linguistic units, a challenge often referred to as the lack of invariance problem (Ladefoged and Broadbent, 1957). To achieve stable perception, listeners rely on context at multiple levels, including lexical information (Ganong, 1980), phonetic structure (e.g., formant patterns and coarticulation) (Ladefoged and Broadbent, 1957), and prosodic cues (e.g., pitch contours) (Wong and Diehl, 2003).

In ambient natural listening environments, the need for contextual modulation is further amplified. Speech is often degraded or ambiguous due to background noise, competing talkers, or reduced signal quality, requiring listeners to rely on contextual cues to maintain perceptual accuracy (Choi et al., 2018; Nusbaum and Magnuson, 1997; Wong et al., 2004; Zhang and Chen, 2016). Successful speech-in-noise perception, for example, depends on dynamically integrating sensory input with expectations formed from recent acoustic and linguistic context (Davis and Johnsrude, 2007; Mattys et al., 2012), highlighting that speech perception is inherently context-dependent and requires continuous updating of expectations under challenging listening conditions.

A growing body of evidence indicates that speech perception in autism is characterized by differences that are closely tied to context-dependent processing. General auditory processing differences in autism span multiple domains (Alcántara et al., 2012; Bonnel et al., 2010, 2003; Dawson et al., 2004; Heaton et al., 2008; Mottron et al., 2000), contributing to variability in how speech signals are encoded and interpreted. At the neural level, representations of basic speech elements (e.g., phonemes or syllables) have been described as less precise (Patel et al., 2022a; Russo et al., 2009, 2008), less robust (Lau et al., 2021; Patel et al., 2022a), and temporally delayed (Chen et al., 2020; Kasai et al., 2005; Lepistö et al., 2007, 2005), suggesting reduced fidelity in encoding the speech acoustic signal itself.

Critically, many of the most clinically salient speech perception differences in autism are not limited to simplified laboratory conditions, but are more prominently documented in dynamic, real-world listening environments where successful perception depends on the use of contextual information. Speech-in-noise (SiN) perception is a prominent example, placing strong demands on listeners to maintain stable representations despite degraded input by drawing on prior acoustic context and expectations. Autistic individuals frequently show difficulty in SiN perception (Alcántara et al., 2012, 2004; Callejo and Boets, 2023; Ruiz Callejo et al., 2023; Smith and Bennetto, 2007), consistent with the idea that reduced contextual modulation may compromise the interpretation of degraded speech.

Converging evidence from other domains further supports context-dependent processing differences in autism. Multispeaker listening paradigms have shown reduced sensitivity to salient speech signals in complex auditory scenes, suggesting differences in how contextual information guides auditory selection (Schwartz et al., 2020a, 2020b). Similarly, reduced sensitivity to prosodic cues such as pitch and rhythm has been reported, particularly when interpretation requires integrating information over broader temporal or linguistic context (Diehl et al., 2008; Järvinen-Pasley et al., 2008; Patel et al., 2023; Peppé et al., 2007). Together, these findings indicate that speech perception differences in autism are not only limited to the encoding of local acoustic details, but also extend to the integration of contextual information over time and across changing listening conditions. Yet, the neural mechanisms that support context-dependent speech processing in ASD remain insufficiently understood.

Context-dependent speech perception is supported by interactive neural mechanisms spanning multiple levels of the auditory system (Arnal and Giraud, 2012; Donhauser and Baillet, 2020; Giraud and Poeppel, 2012). Within predictive processing frameworks, the brain continuously generates expectations from statistical regularities in prior auditory input and uses these predictions to fine-tune sensory encoding in real time (Friston, 2008, 2005; Friston and Kiebel, 2009), with feedback projections enabling higher-order regions to modulate early sensory representations, including those in the auditory brainstem (Chandrasekaran et al., 2014; Chandrasekaran and Kraus, 2010; Skoe et al., 2014).

The frequency-following response (FFR) is a neurophysiological measure reflecting phase-locked neural activity across the auditory pathway that indexes the fidelity of speech encoding. Importantly, FFRs are not purely stimulus-driven (e.g., Galbraith et al., 1998), but are actively modulated by auditory context (Chandrasekaran et al., 2009; Lau et al., 2019, 2017; Parbery-Clark et al., 2011; Skoe et al., 2014; Slabu et al., 2012; Strait et al., 2011; Xie et al., 2018). Specifically, FFRs are enhanced when stimuli are more predictable, but also attenuated with repeated stimulus presentation (Lau et al., 2019, 2017). These effects reflect the interaction of two complementary mechanisms: feedback-based predictive processes that enhance the encoding of expected input, and local stimulus-specific adaptation (SSA) that attenuates responses with repetition (Chandrasekaran et al., 2014; Lau et al., 2019, 2017). Critically, these mechanisms exert opposing influences, such that predictable contexts may enhance responses despite repeated stimulation, positioning the FFR as a sensitive index of how prior context shapes speech encoding. Cortical measures such as mismatch negativity (MMN) have also been used to examine context-dependent auditory processing in autism, but findings have been mixed and speech-based paradigms have not yielded reliable group differences (Chen et al., 2020; Dunn et al., 2008; Green et al., 2020; Kabil et al., 2023; Ludlow et al., 2014; Roberts et al., 2011; Schwartz et al., 2018; Vlaskamp et al., 2017), potentially reflecting MMN’s relatively indirect relationship to ongoing speech encoding.

By contrast, the FFR provides a powerful window into context-dependent modulation of speech encoding as it unfolds over time. As a reflection of phase-locked neural activity, the FFR preserves fine-grained representations of key acoustic features such as fundamental frequency and temporal structure, enabling precise characterization of how contextual and predictive processes modulate speech encoding. In addition, the FFR is highly reliable within individuals, enabling quantification of stable individual differences in how these processes shape encoding across intermediate stages of the auditory system.

A growing body of work using the FFR has documented atypical neural encoding of speech in autistic individuals, who show reduced pitch tracking, degraded encoding of acoustic structure (Lau et al., 2021; Patel et al., 2022b; Russo et al., 2009, 2008), and reduced response strength and stability across trials (Lau et al., 2021; Patel et al., 2022b), reflecting less precise neural representations of speech signals. However, existing work has largely examined FFRs under relatively static listening conditions, characterized by repeated, predictable stimulus sequences with little contextual variability, leaving open the question of how these neural representations are shaped by prior auditory context.

The current study investigates context-dependent modulation of speech encoding in autism using the FFR, examining how predictable versus less predictable auditory contexts influence the fidelity of speech representations in autistic and non-autistic individuals. We hypothesized that speech encoding in autism would be less modulated by contextual predictability, grounded in Bayesian and predictive processing accounts positing reduced weighting of prior expectations relative to incoming sensory input (Lawson et al., 2017, 2014; Palmer et al., 2017; Sinha et al., 2014). While these frameworks are typically formulated as domain-general accounts, with empirical support largely coming from visual tasks (Lawson et al., 2014; Palmer et al., 2017; Pellicano and Burr, 2012), the present study extends this work to auditory speech processing. We predicted that non-autistic individuals would show enhanced neural encoding in more predictable contexts, replicating prior findings, whereas autistic participants would show reduced context-dependent modulation. Beyond group differences, we examined brain–behavior relationships with Speech-in-Noise perception and autism symptom severity, linking neural measures to clinically meaningful variability in auditory and communicative function.

## Methods

### Participants

Participants consisted of 20 autistic individuals (AU group) and 19 non-autistic individuals (non-AU). Participant demographics are presented in Table 1. There were no significant group differences in age (*t*(36.8) = 0.75, *p* = .459). IQ was marginally higher in the non-AU group (*t*(30.7) = −1.88, *p* = .070). A significant difference in sex ratio was observed (χ²(1) = 4.32, *p* = .038), with a higher proportion of males in the AU group (15:5) and females in the non-AU group (7:12).

**Table 1.**
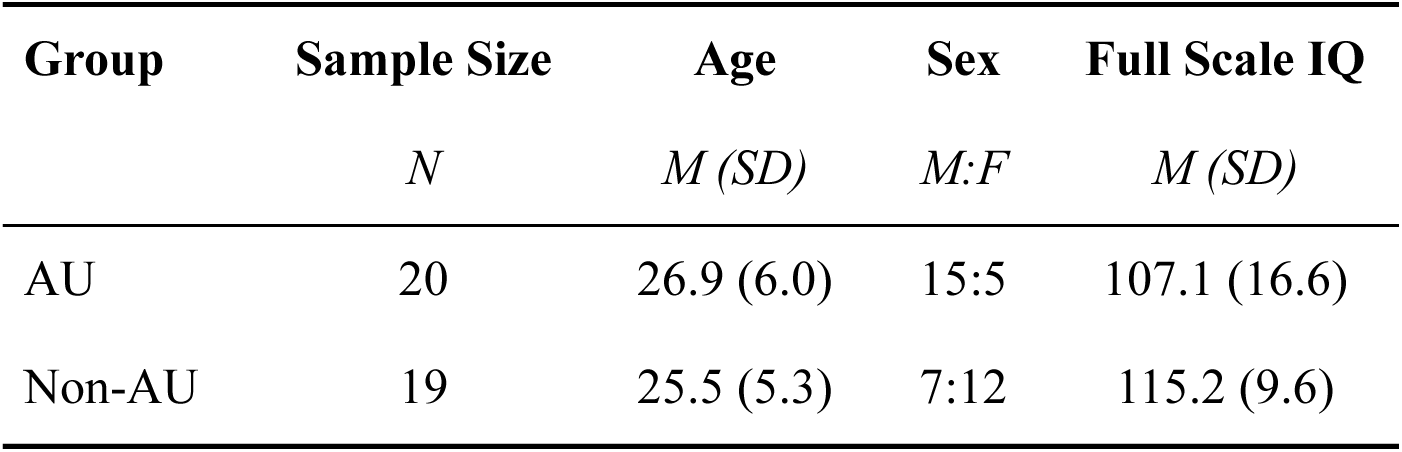
Participant Demographics.

| **Group** | **Sample Size** | **Age** | **Sex** | **Full Scale IQ** |
| --- | --- | --- | --- | --- |
|  | *N* | *M (SD)* | *M:F* | *M (SD)* |
| AU | 20 | 26.9 (6.0) | 15:5 | 107.1 (16.6) |
| Non-AU | 19 | 25.5 (5.3) | 7:12 | 115.2 (9.6) |

All participants were native English speakers with an IQ ≥ 80. All reported no history of dyslexia or neurologic injury or disease. AU group participants had a confirmed clinical diagnosis of autism prior to enrollment, with no co-occurring genetic diagnosis associated with autism (e.g., Fragile X Syndrome). Normal hearing was confirmed by pure-tone air conduction thresholds ≤ 25 dB at 500, 1,000, 2,000, and 4,000 Hz.

All study procedures were approved by the Institutional Review Board of Northwestern University [STU00221550]. All participants provided written informed consent prior to participation, in accordance with the Declaration of Helsinki.

### Clinical-behavioral Measures

IQ was assessed using the Wechsler Abbreviated Scale of Intelligence (WASI) (Wechsler, 1999). Autism symptomatology was characterized in both groups using the Autism Diagnostic Observation Schedule, Second Edition (ADOS-2) (Lord et al., 2012), a gold-standard semi-structured diagnostic instrument for autism.

### Speech-in-noise Measure

Speech-in-noise (SiN) performance was assessed using the Coordinate Response Measure (CRM) task (Bolia et al., 2000). On each trial, two target words (a color and a number) were presented binaurally over headphones in the presence of speech-shaped noise at nine signal-to-noise ratio (SNR) levels ranging from +12 dB to −12 dB in 3-dB increments (Brungart, 2001; Brungart et al., 2001). Two trials were presented per SNR level. Participants reported the perceived color and number using mouse-controlled on-screen sliders. The 50% SNR threshold, taken as the index of SiN performance, was calculated as SiN = 13.5 − .75(r), where r is the total number of target words correctly identified (Tillman and Olsen, 1973).

### Frequency Following Response Experiment Stimuli

Speech stimuli (Fig. 1A) consisted of three intonational contours (*rising*, *level*, and *falling*), with F0 ranges of 179-214 Hz, 188-197 Hz, and 161-207 Hz, respectively, corresponding to interrogative, neutral, and declarative meanings (Sostarics and Cole, 2023). Using Praat, the F0 contours were superimposed onto a naturally produced syllable “ya” (175ms duration and 74 dB intensity) and resynthesized using the overlap-add method.

**Fig. 1.**
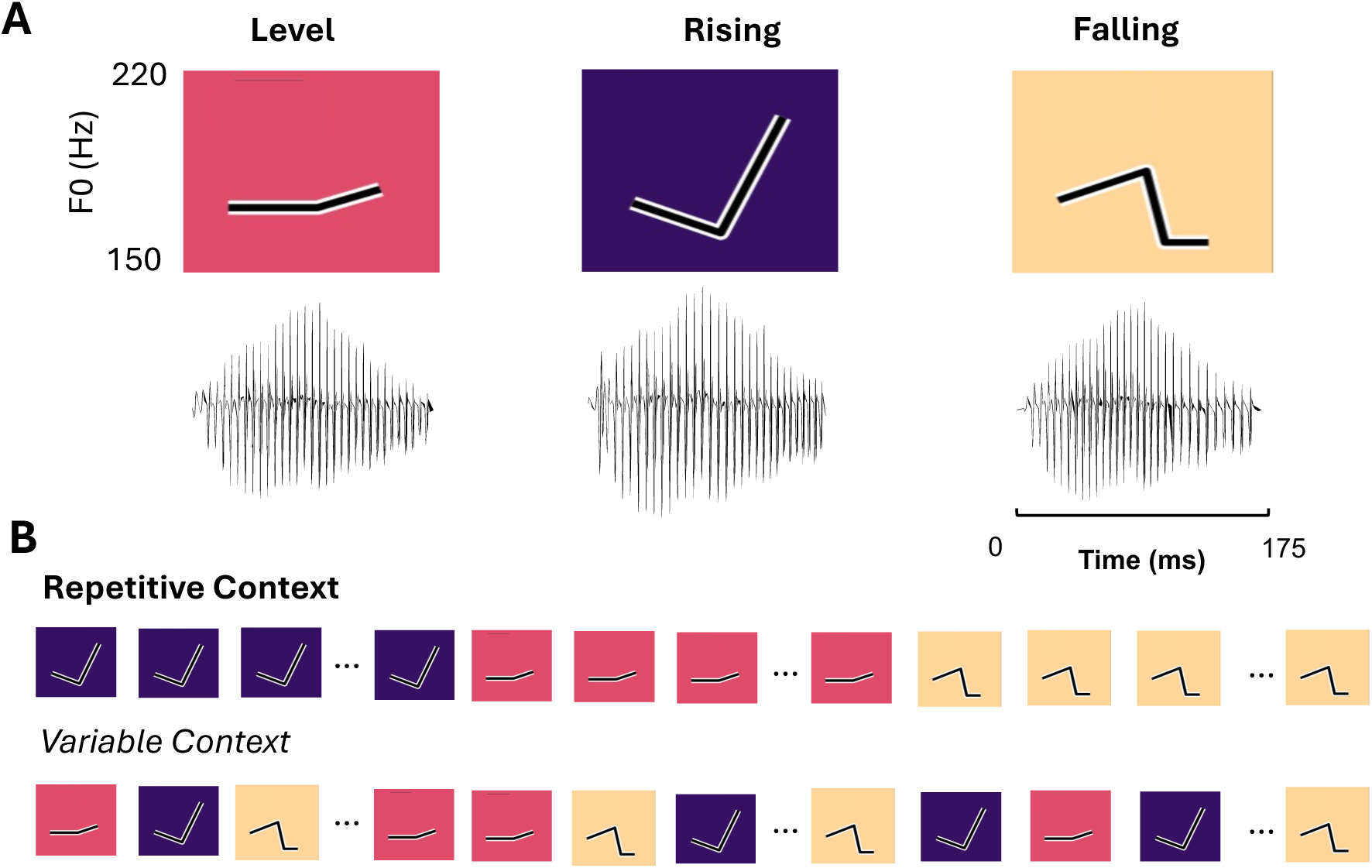
Stimuli and task design. **(A)** Acoustic waveforms and fundamental frequency (F0) contours of the auditory stimuli. Stimuli consisted of 175 ms syllables (“ya”) embedding one of three pitch contours: level, rising, or falling. **(B)** Stimuli were presented in either repetitive or variable contexts. In the repetitive context (top), stimuli were organized into blocks in which a single contour was repeated, with block order randomized across participants. In the variable context (bottom), the three contours were presented in a randomized sequence. Across both contexts, each contour was presented an equal number of times. **Note.** Graphic elements of this figure were adapted from Sostarics and Cole (2023), used with permission.

### Presentation contexts

The three stimuli were presented in two contexts: a *variable* context and a *repetitive* context (Fig. 1B). In the variable context, each stimulus was presented for 1,000 sweeps (500 rarefaction and 500 condensation polarity sweeps) in pseudorandom order, with equal transitional probabilities (33% per stimulus). In the repetitive context, each stimulus was presented repetitively for 1,000 sweeps (in alternating polarities) within a block; the order of the three sets of sweeps was randomized across participants. The order of presentation contexts was counterbalanced across participants. All stimuli were presented with an inter-stimulus interval (ISI) that jittered between 74 ms and 114 ms. The experiment lasted approximately 40 minutes and was administered alongside additional experiments not reported here.

### EEG recording

Stimuli were presented binaurally through electromagnetically shielded insert earphones (ER-3C, Etymotic Research) at 75 dB SPL using the software *Presentation* (Neurobehavioral Systems). Participants were seated in a reclining chair and watched a silent movie during recording, consistent with standard FFR protocols (Chandrasekaran and Kraus, 2010; Skoe and Kraus, 2010). Electrophysiological data were acquired using a BioSemi ActiveTwo system with 32 Ag/AgCl electrodes arranged according to the International 10–20 system, plus electrodes over the left and right mastoids (M1, M2). Voltage offsets were maintained below 35 mV. Data were digitized at 16,384 Hz and band-pass filtered between 0.01 and 100 Hz during acquisition.

### Electrophysiological data preprocessing

Data were preprocessed offline using EEGLAB (Delorme and Makeig, 2004) with the ERPLAB plug-in (Lopez-Calderon and Luck, 2014) in MATLAB (MathWorks). Analyses focused on responses at Cz, re-referenced to the bilateral mastoids (M1, M2), following standard FFR montage conventions (Krizman and Kraus, 2019; Skoe and Kraus, 2010). Data were band-pass filtered from 80 to 2500 Hz (12 dB/octave) and epoched trial-by-trial. Epochs exceeding ±35 µV peak-to-peak were rejected. Responses were averaged across the first 1,000 non-rejected trials per polarity separately for each contour, using a 275-ms epoch comprising a 50-ms pre-stimulus baseline, 175-ms stimulus window, and 50-ms post-stimulus period.

### FFR metrics

Five established FFR metrics were extracted to index complementary aspects of neural encoding (Krizman and Kraus, 2019; Skoe and Kraus, 2010): *neural lag* (ms; encoding latency), *peak autocorrelation* (response periodicity and phase locking), *F0 error* (Hz; pitch encoding accuracy), *signal-to-noise ratio* (magnitude of stimulus-locked neural activity), and *response consistency* (trial-to-trial stability of FFR morphology). Full computational details are provided in the *Supplementary Materials*. All metrics were computed using customized scripts derived from functions from the Brainstem Toolbox (Skoe and Kraus, 2010) on MATLAB.

### Data Availability

Numeric dataset and analytic scripts are available from the OSF database (https://osf.io/9s86h/).

### Statistical Analyses

#### Principal Component Analysis (PCA)

To capture shared variance across multiple FFR metrics indexing complementary aspects of neural encoding (e.g., fidelity, pitch tracking, and timing), we applied principal component analysis (PCA) to derive composite dimensions of neural response variability from the five neural measures: *response consistency*, *peak autocorrelation*, *F0 error*, *neural lag*, and signal-to-noise ratio (*SNR*). Data were first aggregated at the level of participant × group × context × contour, and variables were z-scored prior to PCA. PCA was performed using singular value decomposition. The PCA results, including component loadings and variance explained, are summarized in Table 2.

**Fig. 2.**
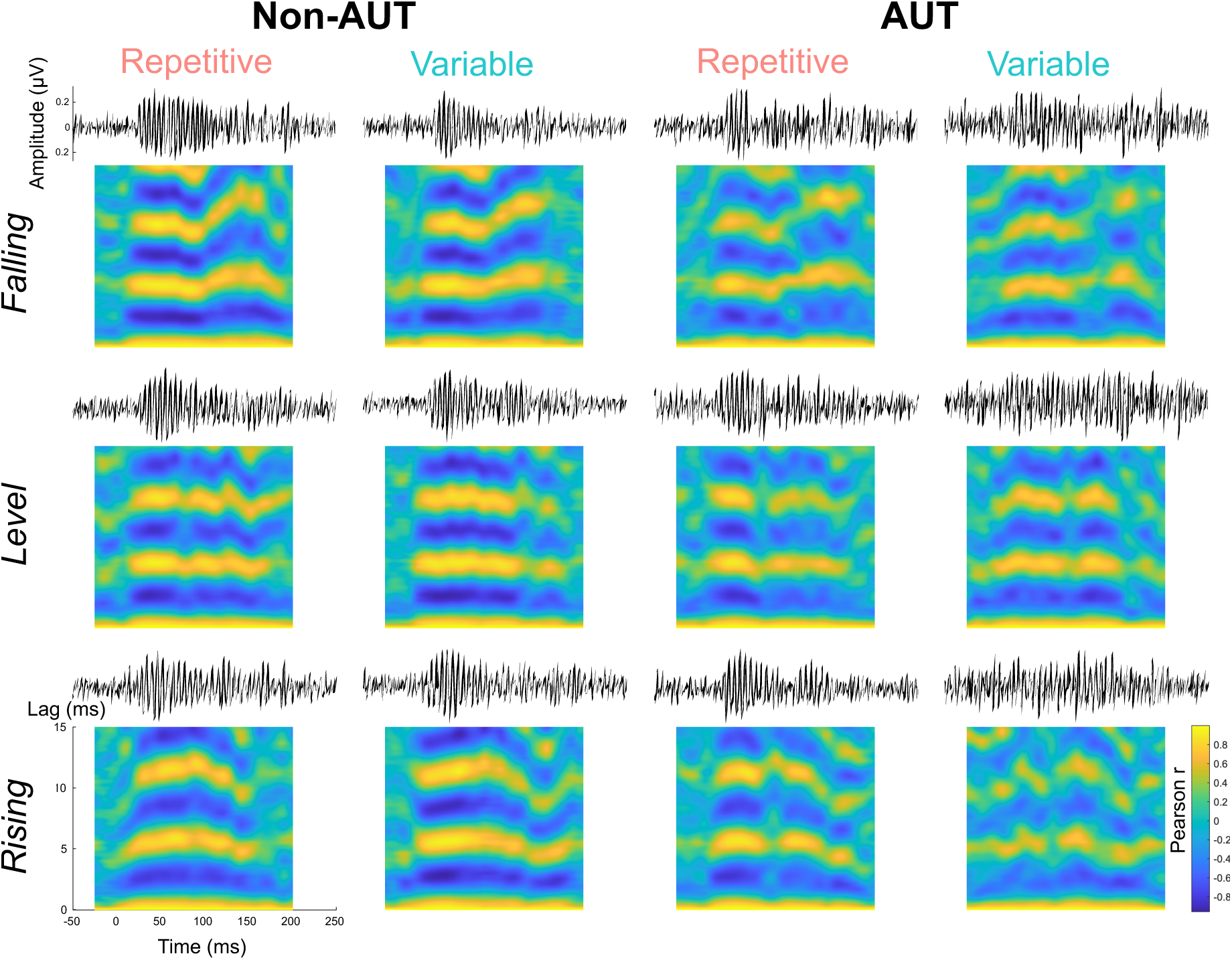
Grand-average frequency-following response (FFR) waveforms (top) and autocorrelograms (bottom) as a function of diagnostic group (AUT vs. Non-AUT), stimulus context (repetitive, variable), and intonation contour (falling, level, rising).

**Table 2:** Principal Component Analysis (PCA) Loadings for Frequency Following Response Metrics.

| **Measure** | **PC1** | **PC2** | PC3 | PC4 | PC5 |
| --- | --- | --- | --- | --- | --- |
| Response Consistency | -0.524 | -0.071 | -0.25 | 0.493 | -0.643 |
| Peak Autocorrelation | -0.533 | -0.113 | -0.04 | 0.372 | 0.75 |
| F0 Error | 0.429 | -0.195 | -0.853 | 0.175 | 0.139 |
| Neural Lag | -0.086 | 0.962 | -0.25 | 0.007 | 0.066 |
| Signal-to-noise Ratio | -0.499 | -0.137 | -0.38 | -0.766 | -0.017 |

Principal components (PCs) retained for interpretation are bolded. Loadings represent correlations between observed variables and principal components.

The first two principal components (PC1 and PC2), accounting for 58.55% and 20.36% of the variance, respectively (cumulative = 78.91%), were retained for analysis, supported by inspection of the scree plot (Fig 3A), which indicated an inflection after the second component.

**Fig. 3.**
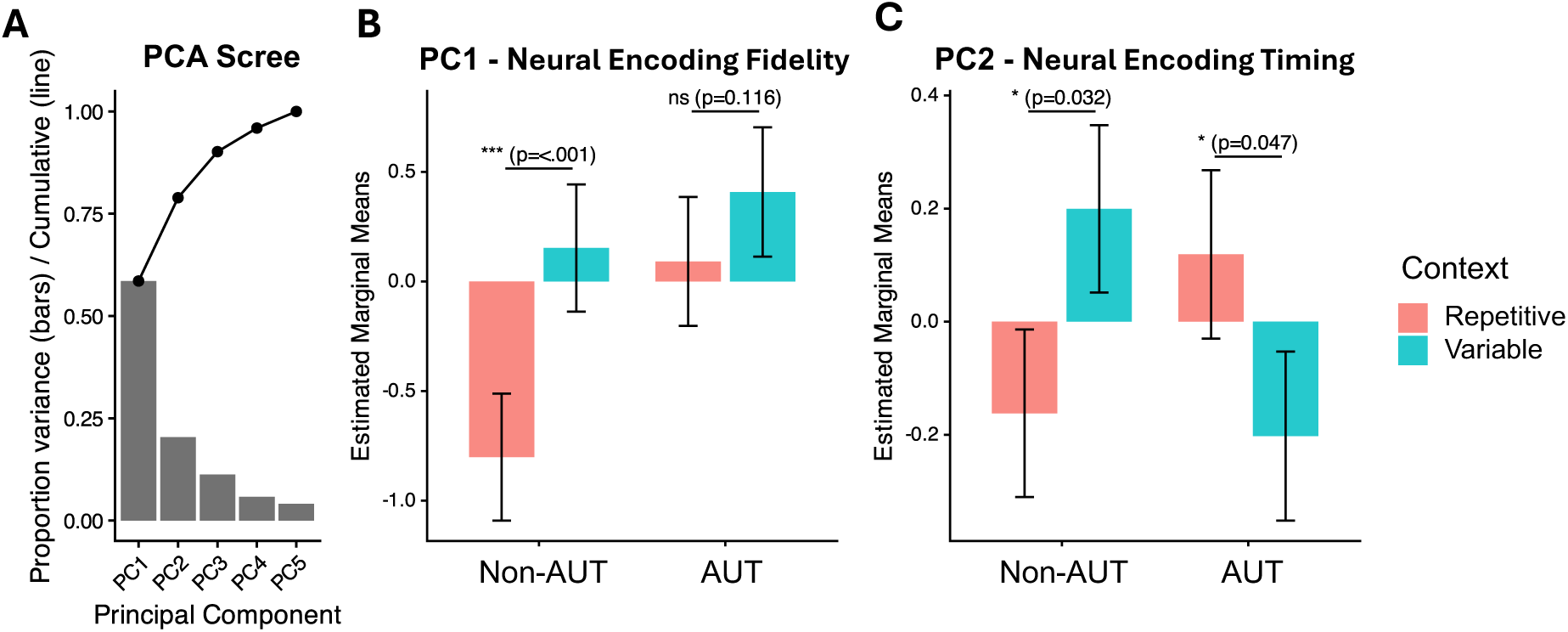
Principal component structure and context-dependent neural encoding across groups. **(A)** Scree plot showing the proportion of variance explained by each principal component (bars) and cumulative variance (line). **(B)** Estimated marginal means (± SE) of the first principal component (PC1), indexing neural encoding fidelity, across repetitive and variable contexts for non-autistic (Non-AUT) and autistic (AUT) groups, averaged across FFRs to all three intonation contour stimuli. The Non-AUT group exhibited higher neural encoding fidelity (more negative PC1 values) in the *repetitive* compared to the *variable* context, whereas no significant context-related differences were observed in the AUT group. **(C)** Estimated marginal means (± SE) of the second principal component (PC2), indexing neural encoding timing, across contexts for both groups. The Non-AUT group exhibited sharper neural encoding timing (more negative PC2 values) in the *repetitive* compared to the *variable* context, whereas the AUT group showed delayed neural encoding timing (more positive PC2 values) in the *repetitive* relative to the *variable* context. **Note:** Pairwise comparisons were conducted between contexts within each group based on estimated marginal means (EMMs); *p < .05; ***p < .001; ns = not significant.

Based on the pattern of loadings, **PC1 was interpreted as reflecting overall neural encoding fidelity**, with higher scores indicating weaker and less consistent responses (i.e., lower response consistency, pitch strength, and SNR, alongside greater F0 error). **PC2 was interpreted as reflecting neural encoding timing**, driven primarily by neural lag.

### Linear Mixed-Effects Models

Separate linear mixed-effects models (LMMs) were fit for PC1 (neural encoding fidelity) and PC2 (neural encoding timing) as dependent variables. Fixed effects included group (AU vs. non-AU), context (repetitive vs. variable), and their two-way interaction, along with covariates for contour, sex, age (z-scored), and IQ (z-scored). Categorical predictors were treatment-coded, with non-AU as the reference level for group, rising contour as the reference level for contour, and the repetitive condition as the reference level for context. Random intercepts were included for participant. FFR counterbalancing order was initially specified as a random effect but was included as a fixed effect in cases of singular fit. Models were estimated using maximum likelihood, and significance of fixed effects was evaluated using Satterthwaite approximations for degrees of freedom.

To examine the effect of context within each group, planned comparisons were conducted using estimated marginal means (EMMs). Within-group contrasts comparing variable versus repetitive contexts were evaluated using *t*-tests with Kenward–Roger approximations for degrees of freedom and Bonferroni correction for multiple comparisons.

### Correlation Analyses

To examine associations between neural context effects and behavioral measures, difference scores (Δ = variable − repetitive) were computed for each participant and principal component. Correlations were conducted within each diagnostic group. Associations with Speech-in-Noise performance were assessed using Pearson correlations, whereas associations with autism symptom severity were assessed using Spearman rank correlations due to the ordinal and potentially non-normal distribution of ADOS-2 calibrated severity scores. ADOS-2 measures included Social Affect (SA), Restricted and Repetitive Behaviors (RRB), and overall severity. P-values were adjusted using the false discovery rate (FDR) procedure separately within each group.

## Results

### Group differences in context-related modulation of neural encoding fidelity (PC1)

To examine group differences in neural encoding fidelity (Fig. 3B), we fit a linear mixed-effects model on PC1 (see Table 3). There was a significant main effect of group, with AU participants showing higher PC1 values than non-AU participants (*b* = 0.892, *t*(49.66) = 2.227, *p* = .031), indicating reduced neural encoding fidelity in the AU group across conditions. There was also a significant main effect of context, with the variable condition associated with higher PC1 values relative to the repetitive condition (*b* = 0.954, *t*(197.29) = 4.537, *p* < .001).

**Table 3.**
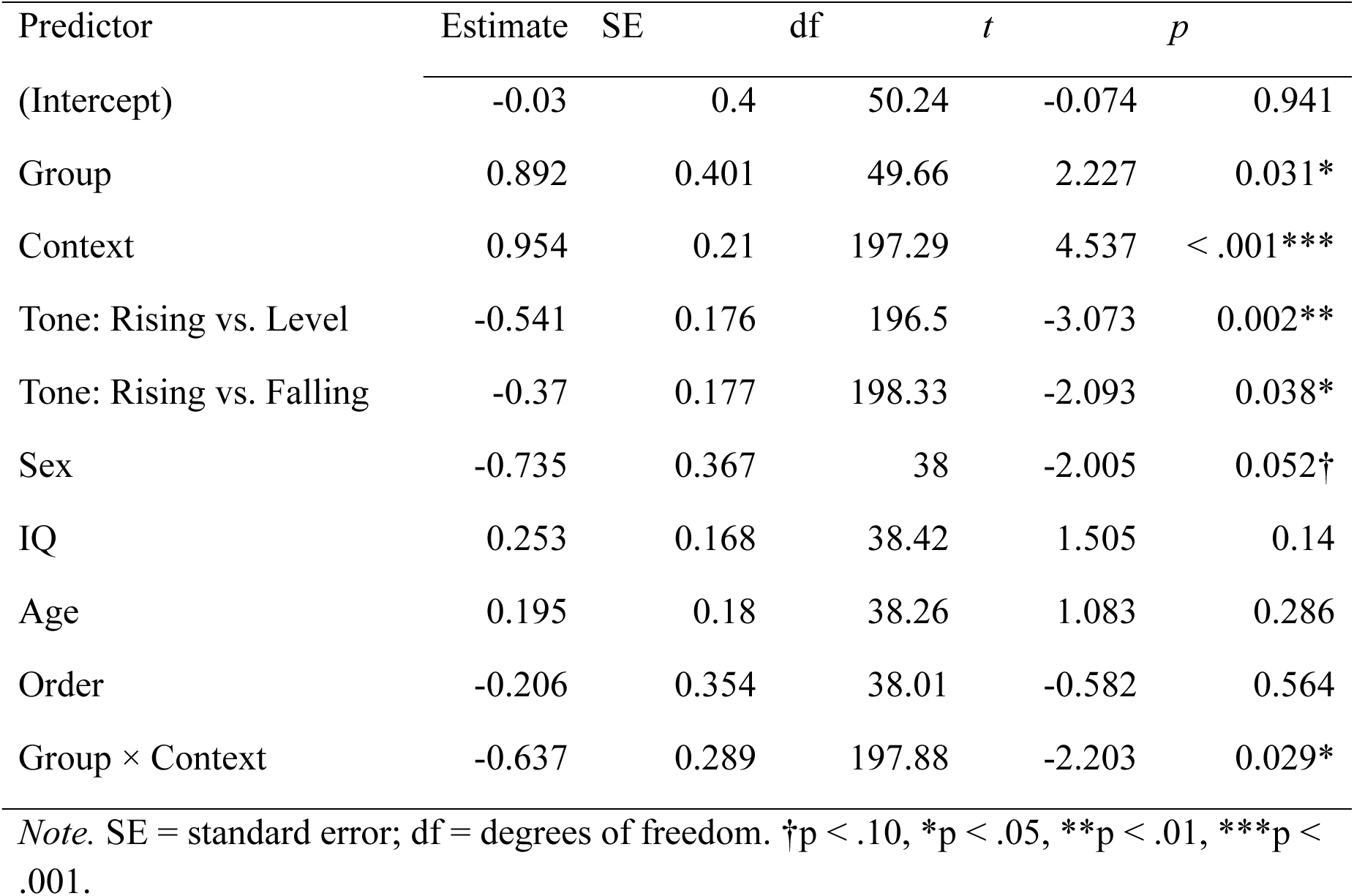
Linear Mixed-Effects Model Results – Principal Component 1 (neural encoding fidelity)

This effect was qualified by a significant group × context interaction (*b* = −0.637, *t*(197.88) = −2.203, *p* = .029), indicating that the magnitude of the context effect differed across groups.

Follow-up comparisons revealed that non-AU participants showed reduced neural encoding fidelity in the variable relative to repetitive condition (Δ = 0.954, *t*(202) = 4.491, *p* < .001), whereas this effect was not significant in the AU group (Δ = 0.317, *t*(203) = 1.577, *p* = .116).

There were also significant effects of contour, with the level pitch contour associated with enhanced neural encoding fidelity relative to the rising contour (*b* = −0.541, *t*(196.50) = −3.073, *p* = .002), and the falling pitch contour also showing enhanced fidelity relative to rising (*b* = −0.370, *t*(198.33) = −2.093, *p* = .038). A marginal effect of sex was observed (*b* = −0.735, *t*(38.00) = −2.005, *p* = .052). No other covariates were significant (all *p*s > .14).

### Group differences in context-related modulation of neural encoding timing (PC2)

To examine group differences in neural encoding timing (Fig. 3C), we analyzed PC2 (a composite index of neural encoding timing, with higher values indicating greater neural lag, see Table 4). The main effect of group was not significant, with AU participants showing numerically higher PC2 values than non-AU participants (*b* = 0.281, *t*(76.85) = 1.372, *p* = .174), indicating no reliable group difference in neural encoding timing across conditions.

**Table 4.**
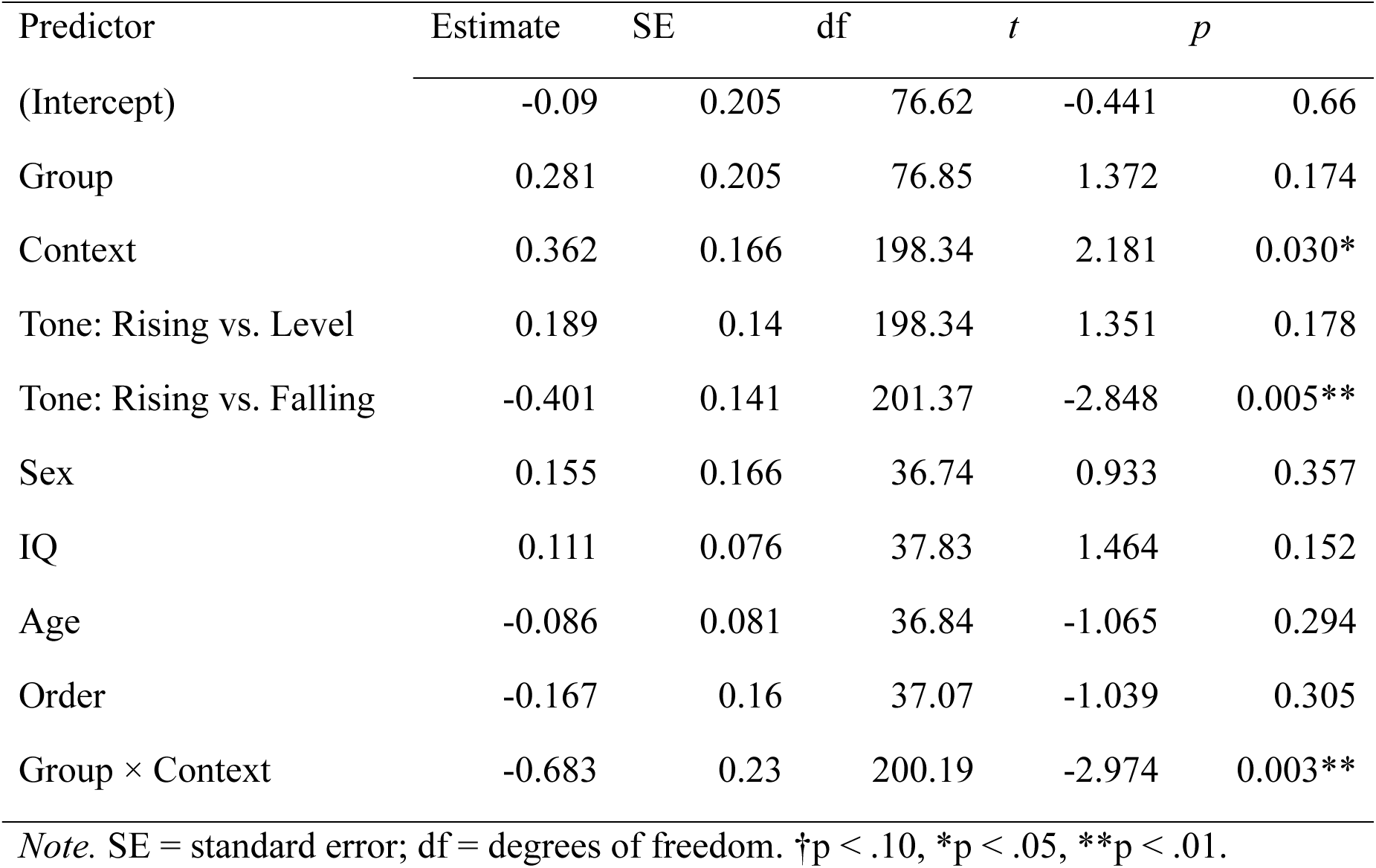
Linear Mixed-Effects Model Results – Principal Component 2 (neural encoding timing)

| Predictor | Estimate | SE | df | *t* | *p* |
| --- | --- | --- | --- | --- | --- |
| (Intercept) | -0.09 | 0.205 | 76.62 | -0.441 | 0.66 |
| Group | 0.281 | 0.205 | 76.85 | 1.372 | 0.174 |
| Context | 0.362 | 0.166 | 198.34 | 2.181 | 0.030* |
| Tone: Rising vs. Level | 0.189 | 0.14 | 198.34 | 1.351 | 0.178 |
| Tone: Rising vs. Falling | -0.401 | 0.141 | 201.37 | -2.848 | 0.005** |
| Sex | 0.155 | 0.166 | 36.74 | 0.933 | 0.357 |
| IQ | 0.111 | 0.076 | 37.83 | 1.464 | 0.152 |
| Age | -0.086 | 0.081 | 36.84 | -1.065 | 0.294 |
| Order | -0.167 | 0.16 | 37.07 | -1.039 | 0.305 |
| Group × Context | -0.683 | 0.23 | 200.19 | -2.974 | 0.003** |

*Note.* SE = standard error; df = degrees of freedom. †*p* < .10, \**p* < .05, \*\**p* < .01.

The main effect of context was significant, with the variable condition associated with higher PC2 values (i.e., longer neural encoding timing / increased neural lag) relative to the repetitive condition (*b* = 0.362, *t*(198.34) = 2.181, *p* = .030).

Importantly, this effect was moderated by group, as evidenced by a significant group × context interaction (*b* = −0.683, *t*(200.19) = −2.974, *p* = .003), indicating that context-related changes in neural timing differed across groups.

Follow-up comparisons showed divergent patterns across groups: non-AU participants exhibited longer neural encoding timing (greater neural lag) in the variable relative to repetitive condition (Δ = 0.362, *t*(204) = 2.160, *p* = .032), whereas AU participants showed shorter neural encoding timing (reduced neural lag) in the variable condition (Δ = −0.321, *t*(208) = −2.002, *p* = .047).

There was also a significant effect of contour, with the falling pitch contour associated with shorter neural encoding timing (reduced lag) relative to the rising contour (*b* = −0.401, *t*(201.37) = −2.848, *p* = .005). Other covariates were not significant (all *p*s > .10).

### Correlation Analyses

In the AU group (Table 5), greater context-related changes in neural encoding fidelity (ΔPC1, defined as the difference between PC1 scores in the variable and repetitive conditions) were associated with stronger social-communication abilities as indexed by ADOS-2 Social Affect scores (ρ = −.61, *p* = .037, FDR-corrected) (Fig. 4A), but poorer speech-in-noise performance (*r* = .58, *p* = .037, FDR-corrected) (Fig. 4B).

**Fig. 4.**
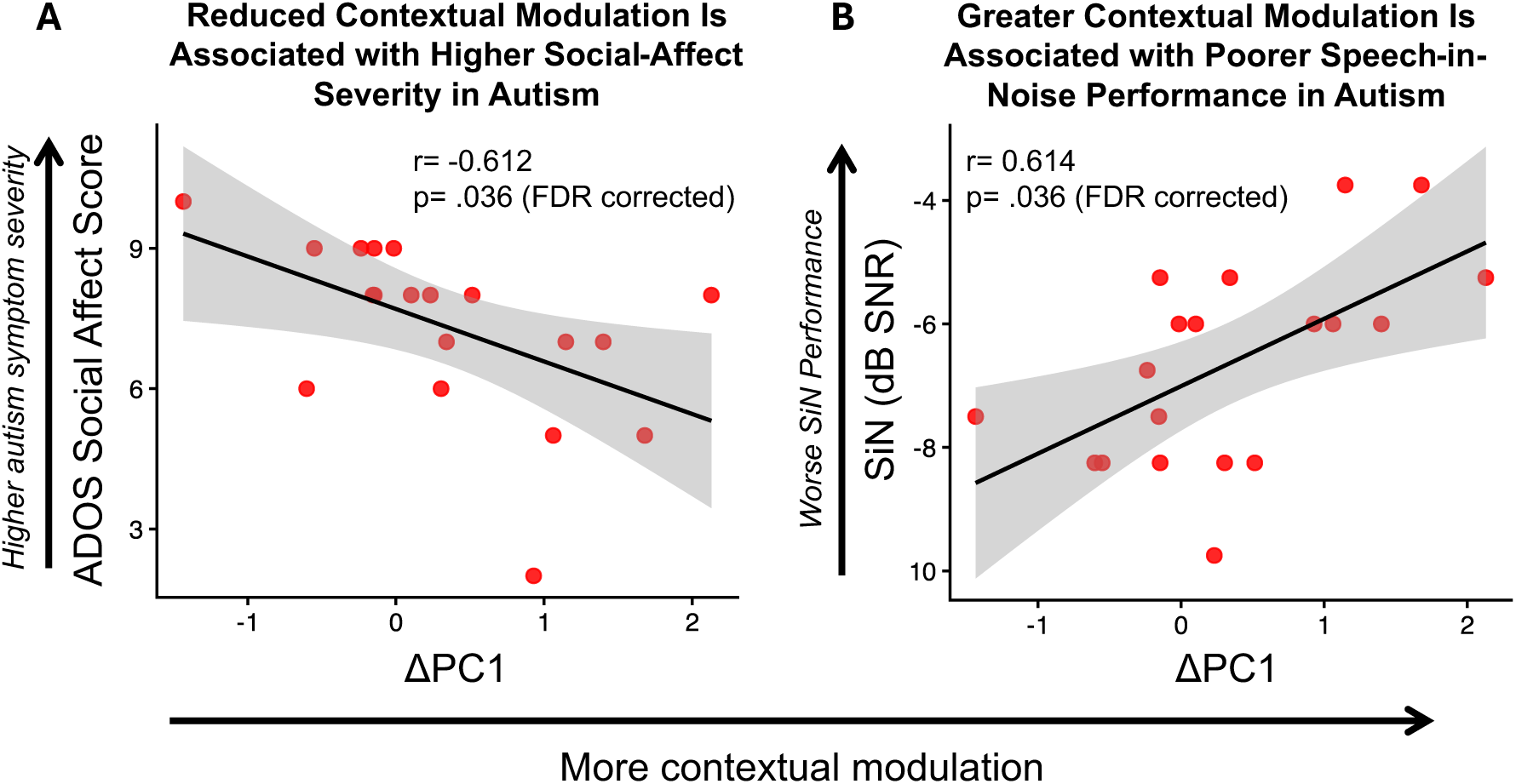
Associations between neural contextual modulation and behavioral measures in the autism group. **(A)** Reduced contextual modulation is associated with higher social-affect severity, indexed by ADOS Social Affect calibrated severity scores. Contextual modulation (ΔPC1) reflects the difference in neural encoding fidelity between the variable and repetitive contexts, calculated as the value in the variable context minus the value in the repetitive context, with more positive values indicating greater modulation. **(B)** Greater context-dependent modulation of neural encoding is associated with poorer speech-in-noise performance, indexed by lower signal-to-noise ratio (SNR) thresholds (dB; lower values indicate better performance). P-values are false discovery rate (FDR) corrected.

**Table 5:** Correlations Between Degrees of Contextual Modulation (ΔPC1, ΔPC2) and Clinical-behavioral Measures in the AUT group.

| Component | Measure | $r$ | $p$ | p (FDR) |
| --- | --- | --- | --- | --- |
| $\Delta PC1$ | ADOS-2 Restricted and Repetitive Behaviors | 0.239 | 0.356 | 0.407 |
|  | ADOS-2 Social Affect | -0.612 | 0.009** | 0.036* |
|  | ADOS-2 Overall | -0.368 | 0.147 | 0.293 |
|  | SiN | 0.614 | 0.009** | 0.036* |
| $\Delta PC2$ | ADOS-2 Restricted and Repetitive Behaviors | -0.296 | 0.248 | 0.331 |
|  | ADOS-2 Social Affect | -0.382 | 0.131 | 0.293 |
|  | ADOS-2 Overall | -0.301 | 0.240 | 0.331 |
|  | SiN | -0.102 | 0.697 | 0.697 |

*Note.* $r$ = Pearson correlation coefficient. FDR = false discovery rate correction within analysis. \* $p < .05$ , \*\* $p < .01$ .

No significant associations were observed between ΔPC1 and ADOS-2 Restricted and Repetitive Behaviors or overall severity, or between ΔPC2 and speech-in-noise performance or any ADOS-2 domain score (all FDR-corrected ps > .244).

In the non-AU group (Table 6), no significant associations were observed between neural context effects (ΔPC1 or ΔPC2) and Speech-in-Noise performance or any ADOS-2 domain score (all FDR-corrected *p*s > .774), suggesting that relationships between neural context effects and phenotypic variability may be specific to the autistic group.

**Table 6:**
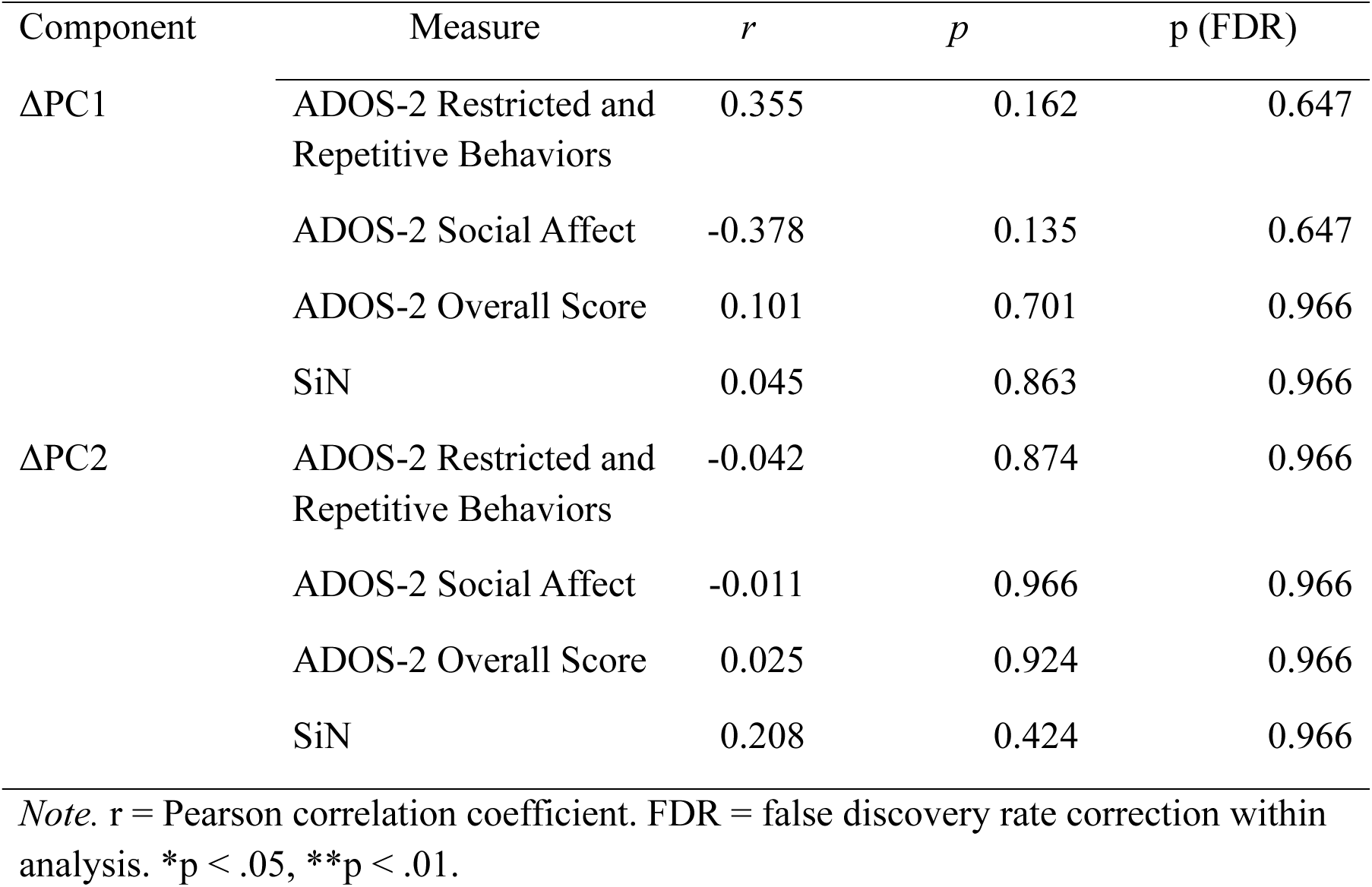
Correlations Between Degrees of Contextual Modulation (ΔPC1, ΔPC2) and Clinical-behavioral Measures in the Non-AUT group.

| Component | Measure | $r$ | $p$ | p (FDR) |
| --- | --- | --- | --- | --- |
| $\Delta PC1$ | ADOS-2 Restricted and Repetitive Behaviors | 0.355 | 0.162 | 0.647 |
|  | ADOS-2 Social Affect | -0.378 | 0.135 | 0.647 |
|  | ADOS-2 Overall Score | 0.101 | 0.701 | 0.966 |
|  | SiN | 0.045 | 0.863 | 0.966 |
| $\Delta PC2$ | ADOS-2 Restricted and Repetitive Behaviors | -0.042 | 0.874 | 0.966 |
|  | ADOS-2 Social Affect | -0.011 | 0.966 | 0.966 |
|  | ADOS-2 Overall Score | 0.025 | 0.924 | 0.966 |
|  | SiN | 0.208 | 0.424 | 0.966 |

*Note.* $r$ = Pearson correlation coefficient. FDR = false discovery rate correction within analysis. \* $p < .05$ , \*\* $p < .01$ .

## Discussion

The present study investigated context-dependent modulation of speech encoding in autism by examining speech-evoked frequency-following response (FFR). Consistent with hypotheses, autistic participants showed reduced context-dependent modulation of neural encoding relative to non-autistic participants, particularly in measures indexing encoding fidelity (PC1) and timing (PC2). Individual differences in context-related neural modulation were further associated with speech-in-noise perception and autism symptom severity in social communication, linking these neural effects to clinically relevant variability.

Across conditions, autistic participants showed reduced neural encoding fidelity relative to non-autistic participants, consistent with prior work reporting less precise and less robust neural encoding of speech in autism (Lau et al., 2021; Patel et al., 2022b; Russo et al., 2009, 2008). Importantly, this group difference was modest relative to the context-dependent effects observed, suggesting that alterations in context-dependent modulation may be more pronounced than differences in baseline encoding itself.

Consistent with prior work, non-autistic participants showed enhanced neural encoding in more predictable contexts, replicating findings that speech encoding is strengthened when stimuli support stronger top-down predictions (Chandrasekaran et al., 2009; Lau et al., 2019, 2017; Parbery-Clark et al., 2011; Skoe et al., 2014; Slabu et al., 2012; Strait et al., 2011; Xie et al., 2018). Whereas prior studies have primarily examined segmental contrasts or pitch patterns in lexical tonal contexts (Chandrasekaran et al., 2009; Lau et al., 2019, 2017; Skoe et al., 2014; Xie et al., 2018), the present findings extend this work to intonational pitch patterns in English, suggesting that context-dependent mechanisms generalize across different levels of speech representation.

In contrast to the robust context effect observed in non-autistic participants, results showed that autistic participants did not exhibit reliable modulation of FFRs as a function of contextual predictability. This pattern was most clearly captured by PC1, which indexes the fidelity of phase-locked speech encoding and has shown reliable context-dependent enhancement in prior work in non-autistic individuals (Lau et al., 2019, 2017). This results pattern is consistent with our hypothesis of reduced predictive tuning in autism, in which prior context exerts less influence on shaping ongoing neural representations. Findings also align with Bayesian and predictive processing accounts which posit reduced weighting of prior expectations relative to incoming sensory input in autism (Hadad and Yashar, 2022; Lawson et al., 2014; Palmer et al., 2017), and extends these accounts to early stages of auditory speech encoding.

A complementary pattern emerged in neural encoding timing (PC2), providing further mechanistic insight into context-dependent modulation differences in autism. Autistic participants showed faster neural encoding timing in variable contexts relative to repetitive contexts, a pattern opposite to that observed in non-autistic participants, who showed faster encoding in repetitive contexts.

This dissociation could reflect a shift in the balance between feedback-based predictive processes, mediated by descending corticofugal pathways from auditory cortex to subcortical structures (Chandrasekaran et al., 2014; Suga and Ma, 2003), and local stimulus-specific adaptation (SSA) mechanisms (Chandrasekaran et al., 2014; Lau et al., 2017). Predictive processes enhance encoding of expected input based on prior context, whereas SSA attenuates responses with repeated stimulation. Because these mechanisms exert opposing influences, the observed response reflects their relative balance. In neurotypical listeners, predictive mechanisms dominate, shaping encoding even under repetitive stimulation (Lau et al., 2017). In autism, reduced predictive modulation may allow SSA to exert relatively stronger influence, such that responses in variable contexts (i.e., where adaptation is reduced) become faster, producing the observed reversal. This pattern is consistent with diminished predictive modulation, enhanced local adaptation, or a combination of both.

Meanwhile, prior work suggests that context-dependent modulation of the FFR is sensitive to attentional state. Under high attentional load (e.g., during a demanding visual task), the typical enhancement of neural encoding in predictable contexts can be reduced or even reversed, with relatively stronger encoding observed in more variable contexts (Xie et al., 2018). Differences in attentional allocation and engagement have been reported in autism (Keehn et al., 2013; Robertson and Baron-Cohen, 2017), and although task instructions were identical across groups and attention was directed away from the auditory stimuli, attentional engagement was not directly measured and may have contributed to the observed differences. Future studies incorporating direct measures or manipulations of attention (e.g., eye tracking or task-based attentional load) will be important for clarifying how attentional and predictive processes interact to shape context-dependent speech encoding in autism.

Correlation analyses revealed that greater context-related changes in neural encoding fidelity (ΔPC1) were associated with lower severity in the ADOS-2 Social Affect domain within the AU group, whereas no associations were observed for Restricted and Repetitive Behaviors or overall autism symptom severity. This selective relationship with the Social Affect domain suggests that context-dependent auditory encoding may relate more closely to social-communicative processes than to broader autism symptomatology. Given that successful communication relies on integrating contextual information over time (Bruner, 1990), reduced sensitivity to context at early stages of auditory encoding may co-vary with difficulties interpreting ongoing input in socially meaningful ways, pointing to a meaningful link between early auditory encoding and higher-level social-communicative functioning.

Associations with Speech-in-Noise perception provide more direct insight into the functional relevance of these neural differences. While post-hoc group comparisons revealed poorer SiN performance in the autism group (AU: *M* = −6.60 dB, TD: *M* = −7.70 dB; *t*(37.0) = 2.19, *p* = .035, *d* = 0.70), within the autism group, greater ΔPC1 was associated with poorer SiN performance. This contrasts with prior work in non-autistic samples reporting associations between greater contextual modulation and better SiN performance (Chandrasekaran et al., 2009), suggesting that context-dependent modulation may not operate as a uniformly beneficial mechanism in autism, with its functional impact depending on the reliability of available input.

Speech-in-noise perception places strong demands on contextual integration to support interpretation of degraded input, and autistic individuals frequently show difficulties in such conditions (Alcántara et al., 2012, 2004; Callejo and Boets, 2023; Ruiz Callejo et al., 2023; Smith and Bennetto, 2007). In the CRM task, speech-shaped noise degrades both the target signal and the reliability of contextual information. Given that baseline speech encoding is, on average, less robust in autism (Lau et al., 2021; Patel et al., 2022b; Russo et al., 2009, 2008), stronger neural contextual modulation (ΔPC1) in autistic individuals may therefore reflect increased sensitivity to contextual cues that, when unreliable, amplifies interference and exacerbates perceptual uncertainty rather than improving performance.

In sum, these findings link variability in context-dependent neural speech encoding to conceptually related behavioral differences in autism, pointing toward a brain-based mechanistic account of elements of the autism clinical phenotype. Findings suggest that individuals with stronger neural contextual modulation may rely more heavily on top-down expectations, which could be advantageous when contextual information is stable and informative (e.g., social interaction in the ADOS-2 task), but perhaps less effective when sensory and contextual cues are degraded. Together, these results link variability in contextual modulation to clinically relevant behavioral features of autism, suggesting that the functional impact of contextual modulation depends on input reliability and task demands.

### Limitations and Future Directions

The present findings provide evidence for context-dependent modulation of speech encoding and its behavioral relevance in autism; several considerations are important for interpreting these findings and guiding future research.

First, the correlational nature of the study limits causal inference regarding the mechanisms underlying context-dependent modulation. While the observed patterns are consistent with a shift in the balance between predictive and locally driven processes, these mechanisms were not directly manipulated and their relative contributions remain inferential. Future work combining this paradigm with functional neuroimaging or connectivity-based analyses (Sitek et al., 2022, 2019) may help clarify the neural circuitry supporting these effects.

In addition, the current paradigm employed highly controlled and relatively artificial auditory contexts. Although such designs are critical for isolating the contribution of predictive and adaptation mechanisms to speech encoding, they may not fully capture the complexity of real-world listening environments. Future studies using more ecologically valid stimuli, such as continuous or naturalistic speech, will be important for determining how context-dependent modulation operates under realistic communicative conditions.

The scope of behavioral characterization beyond autism severity in the present study was also relatively focused, targeting Speech-in-Noise perception based on a priori hypotheses regarding its relevance to context-dependent auditory processing. Context-dependent auditory processing may potentially impact other domains of functioning in autism (e.g., language ability, pragmatic communication, and real-world listening function); future work incorporating more comprehensive behavioral measures will help clarify the broader clinical relevance of these differences.

## Conclusion

The present study provides evidence that context-dependent modulation of speech encoding, i.e., the process by which prior sound patterns shape how the brain encodes incoming speech, is altered in autism, affecting both the fidelity and temporal dynamics of neural responses along the auditory pathway. This capacity is fundamental to robust speech perception, particularly in the dynamic, noisy listening environments of everyday life. Contrasting predictable and less predictable auditory contexts, analyses revealed that autistic individuals exhibited reduced sensitivity to contextual structure during speech processing, consistent with a shift in the balance between feedback-based predictive mechanisms and locally driven sensory processes. These neural differences were associated with variability in ADOS-2 Social Affect scores and speech-in-noise perception, linking context-dependent encoding to both social-communicative function and listening under degraded conditions, with effects depending on input reliability and task demands.

Findings also extend theoretical accounts of altered predictive processing in autism, often framed within Bayesian and predictive coding perspectives (Hadad and Yashar, 2022; Lawson et al., 2017, 2014; Palmer et al., 2017; Pellicano and Burr, 2012; Sinha et al., 2014), to the domain of auditory speech perception, providing a neurophysiological framework for understanding how such differences may impact everyday communication in autism. By identifying context-dependent modulation as a potential mechanism linking neural encoding to perceptual function, this work helps to highlight potential targets for future research aimed at characterizing phenotypic variability of autism and developing interventions that support adaptive use of context in complex listening environments.

## Acknowledgments

This research was supported by grants from the National Institutes of Health (R21DC022031 to JL; R01DC010191, R01DC021849, R01DC022484, R01MH091131, and R03DC018644 to ML, and R01DC013315 to BC)

