## Supplementary Materials for "Neurophysiological Evidence for Reduced Use of Prior Sound Patterns to Shape Speech Processing in Autism"

### Frequency Following Response Metrics

*Neural lag* (in ms) was defined as the time shift yielding the maximum positive cross-correlation between the stimulus and FFR waveforms, estimating neural conduction latency and used to temporally align signals for subsequent analyses.

*Peak autocorrelation* quantified response periodicity and neural phase locking. The 175-ms response window (shifted by neural lag) was segmented into 125 overlapping 50-ms bins (49-ms overlap). Within each bin, the waveform was time-shifted in 1-ms steps and Pearson's r computed at each delay; the maximum value per bin was retained, with higher values reflecting greater periodicity. Peak autocorrelation was defined as the mean maximum r across all 125 bins.

*F0 error* (Hz) was computed as the mean absolute distance between the stimulus F0 and response F0 across the 125 time bins, indexing the accuracy of neural pitch encoding.

*Signal-to-noise ratio (SNR)* was computed to assess the magnitude of neural activation. Root mean square (RMS) amplitudes were calculated for both the FFR window (from *neural lag* to *neural lag* + 175 ms) and the pre-stimulus baseline window (−50 ms to *neural lag*). RMS amplitude was defined as the mean absolute voltage across all sample points within each window (µV). SNR was defined as the ratio of FFR to baseline RMS amplitude, with larger values indicating stronger stimulus-locked neural activity relative to background noise.

*Response consistency* quantified trial-to-trial stability of neural encoding using a bootstrapping approach. For each participant and stimulus, epochs from the two polarities were randomly resampled across 300 iterations; on each iteration, half the trials per polarity were averaged to generate two subaverage waveforms, and Pearson's r was computed between them. Response consistency was defined as the mean r across iterations, with higher values indicating greater stability of FFR morphology.
